# Integrated analysis of gene correlation reveals disordered relationship between metabolism and immunity in tumor microenvironment

**DOI:** 10.1101/2020.03.08.982850

**Authors:** Zixi Chen, Jinfen Wei, Yuchen Yuan, Ying Cui, Yanyu Zhang, Yumin Hu, Hongli Du

**Affiliations:** School of Biology and Biological Engineering, South China University of Technology, Guangzhou 510006, China; Sun Yet-Sen University Cancer Center, State Key Laboratory of Oncology in South China, Collaborative Innovation Center for Cancer Medicine, Guangzhou 510060 Guangdong, China

**Keywords:** spearman correlation, metabolism, immunity, pan-cancer, immune checkpoints

## Abstract

**Background:** Metabolism reprogramming and immune evasion are the most fundamental hallmarks for cancer survival. The complex interactions between metabolism and immune systems in tumors and their microenvironment is complicated. Researching on the correlation changes between metabolic and immune related-genes in normal and tumor tissues would help to reveal these complex interactions.

**Methods:** In this study, the mRNA profiles across 11 cancer types was obtained from The Cancer Genome Atlas (TCGA). Then, the spearman’s correlation coefficient was calculated between metabolic and immune related-genes for each sample group.

**Results:** Our results showed that the number of correlated gene pairs was reduced significantly in tumor tissues compared with those of normal tissue, especially in KIRC, KIRP and STAD. Functional enrichment analysis for the universal (the pairs appeared in more than 2 cancer types) and specific (the pairs only in one specific cancer type) gene pairs across cancer types revealed top pathways which appeared in tumor and normal samples, such as phosphatidylinositol signaling system and inositol phosphate metabolism. Thereinto, the pairs in normal tissues missing in tumors may indicate they are important factors affecting immune system, such as, DGKs and PIP4ks. The correlation analysis between immune checkpoint and metabolism genes also showed a reduced correlation in tumor and had the tissue specificity, such as, *FUT8* was strongly correlated with *PDCD1* in the HC of STAD and they had a weaker correlation in other normal tissues and tumor types.

**Conclusions:** Our study provides a novel strategy for investigating interaction of tumor immune and metabolism in microenvironment and offers some key points for exploring new targets including metabolic targets and immunomodulator of immune checkpoints.

## Background

Recently, inflammation and immune evasion are considered as hallmarks of cancer progression, highlighting the direct involvement of immune cells^[1, 2]^. The study on immunology has made great progress in cancer treatment, and the role of immune cells in cancer progression is well-recognized^[3, 4]^. Cancer immunotherapy has subverted the traditional concept of treatment, such as immune checkpoint inhibitors, cancer vaccines and chimeric antigen receptor redirected T (CAR-T) cell therapy^[5-8]^.

The research progresses on cancer has indicated that metabolic reprogramming is another hallmark of cancer^[9]^. Cells that are common to many cancers that do not produce enough energy due to lack of oxygen, carbohydrate or protein use altered metabolic pathways to ensure their survival. Thus, malignant cells acquire the molecular materials and energy necessary to sustain proliferation through unusual metabolic pathways. The tendency of malignant cells to utilize glucose via the process of glycolysis irrespective of the oxygen availability is known as “aerobic glycolysis” which was pointed by Otto Warburg^[10, 11]^.

Though metabolic reprogramming and evasion of immune surveillance are distinct processes, recent evidence has accumulated to show that specific metabolic signatures as an important regulator in both innate and adaptive immunity in human cancer^[12]^. The immune response is associated with dramatic modifications in tumor microenvironment (TME) metabolism, including depletion of nutrients, increased oxygen consumption, and the generation of reactive nitrogen and oxygen intermediates^[13]^. For example, the bile metabolism of bile acids in gut microbiome can influence the NKT cell–driven killing ability^[14]^. Similarly, depletion in amino acid and nutrient depletion were reported for T cells functions on anti-tumor effect^[15]^. Besides, the crosstalk between immune cells and other cells could influence the immunity^[16, 17]^. Taken together, the findings of immune cells function have been showed altered by the TME extensively in cancers.

The metabolism and immunity are fundamentally linked among the malignant cell, immune cell and the microenvironment cells around them. Thus, there is a renewed interest to exploit the link between these two cancer-related processes in order to develop potent anticancer therapeutics. The correlation between genes expression level can reflect the relation between biological processes^[18-20]^. Up to now, some researchers have excavated and studied metabolic^[21, 22]^ and immune^[23, 24]^ related genes separately based on TCGA database, however, few studies has been done on the expression relationship between metabolic and immune related genes in tumor samples. Thus, there is an interest to exploit the link between these two cancer-related processes in order to develop potent anticancer therapeutics. In this study, we analyzed 11 different cancers from the TCGA database, including 5645 cancer and normal samples, describing the co-expression relationship between metabolic and immune genes in pan-cancer. Our findings will provide a comprehensive data basis for further studies of metabolic and immune associations in tumor tissue.

## Methods

### Data obtaining and preprocessing

The mRNA expression data and clinical information data were obtained from The Cancer Genome Atlas (TCGA) program through the NCI’s Genomic Data Commons (GDC) website (https://gdc.cancer.gov/)^[25]^. 5645 samples across 11 cancers were included in this research. Samples of colon adenocarcinoma (COAD) and rectum adenocarcinoma (READ) were mixed into colorectal carcinoma (CRC), thus left 10 cancer types in the following study. According to the clinical information, each cancer was first classified into 5 groups: healthy control (HC) and tumor samples from stage 1 to stage 4. Then stage 1 and stage 2 were merged into early stage, while stage 3 and stage 4 were merged into advanced stage, the sample details was presented in Supplement Table 1.

**Table 1.**
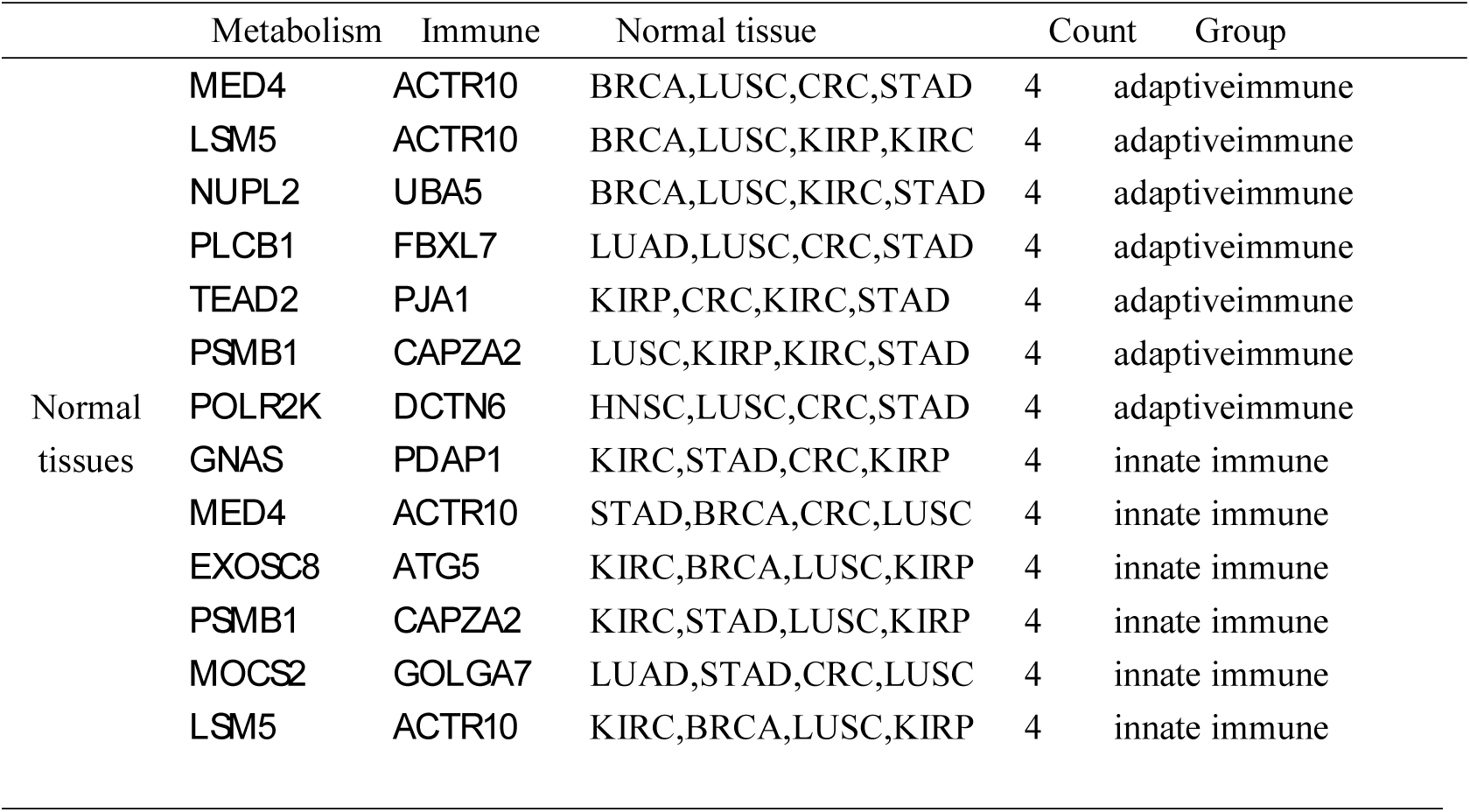
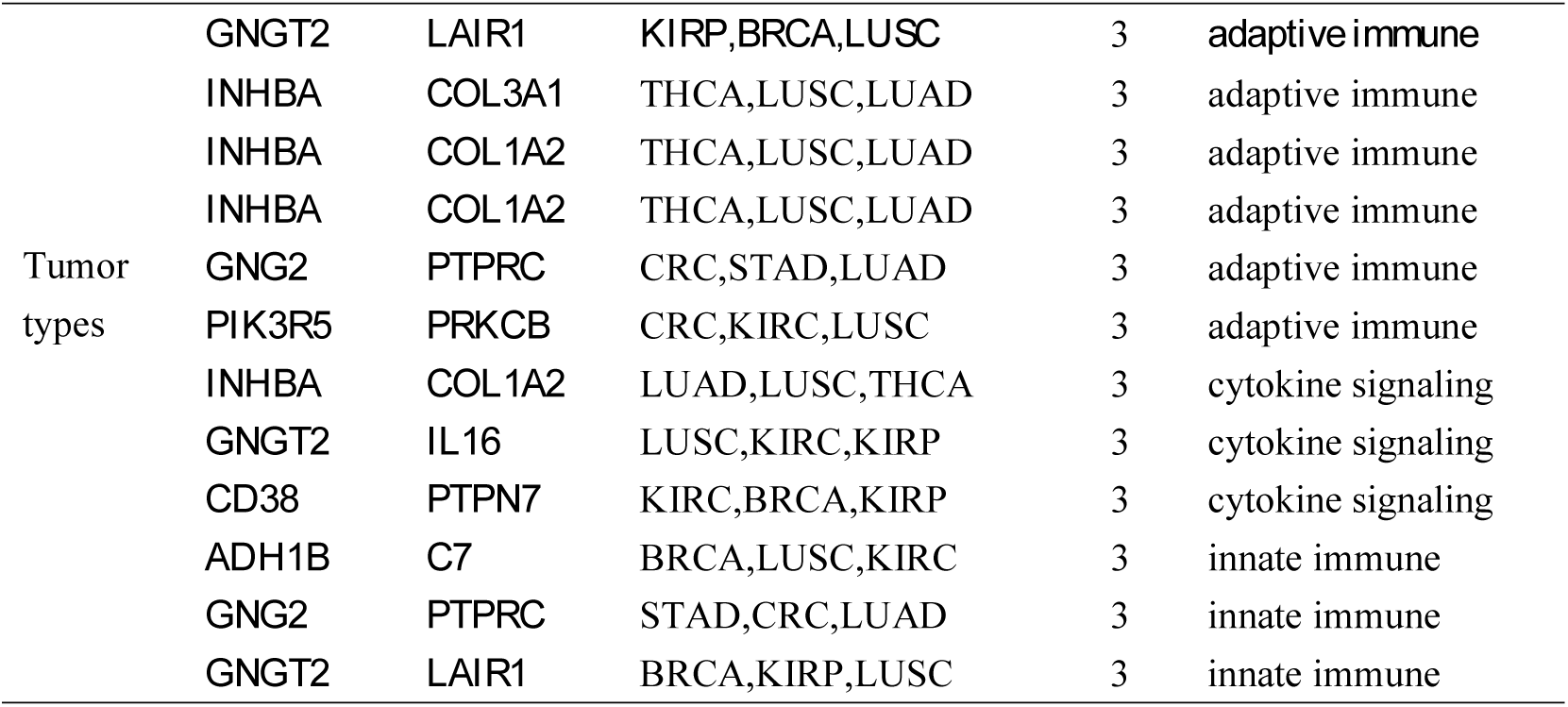
Top universal metabolism-immune gene pairs in HC and tumors.

The list of all human cellular metabolism genes was downloaded from ccmGDB^[26]^. The immune-related gene list was obtained from the Reactome database, these genes were grouped according to three pathways in Reactome Pathway Browser(http://www.reactome.org/): adaptive immunity, innate immunity and cytokine signaling pathway in immune system^[27, 28]^. The genes were listed in Supplement Table 2.

For the RNA-seq data, raw read counts were used to calculate TPM^[29-31]^. Annotation file from GENCODE (https://www.gencodegenes.org/)^[32]^ was used to calculate the length of each gene, which was needed in the calculation of TPM. For each gene, mean TPM was calculated, gene with mean TPM less than 1 was considered as low expressed gene and filtered out.

### Co-expression analysis and DEG analysis

Python (https://www.python.org/) and the “spearmanr” function in package scipy (https://www.scipy.org/) were used to calculate Spearman’s rank correlation in our study^[33]^.

For each cancer type, co-expression analysis was conducted in 7 groups of samples: normal control, tumor from stage 1 to stage 4, early stage tumor, advanced stage tumor. In each sample group, all metabolism genes were paired with 3 groups of immune-related genes: adaptive immunity, innate immunity and cytokine signaling pathway, respectively. There were 21 co-expression analyses to be conducted for each cancer. To calculate Spearman’s *r* value, TPM values of these pairs in each sample were used. Pairs with |*r*| > 0.8 were determined as strongly co-expressed pairs^[34]^. In order to present the landscape when applying different threshold values to our data, the number of co-expressed pairs with |*r*| > 0.7, |*r*| > 0.8 and |*r*| > 0.9 was counted.

In order to clarify the differences between normal and tumor tissues, a further filter was conducted between HC with tumor (early and advanced stage). Pairs with |*r*__hc_| > 0.8 in HC and |*r*__tumor_| < 0.4 were defined as HC specific metabolism-immune gene pairs. Pairs with |*r*__hc_| < 0.4 in HC and |*r*__tumor_| > 0.8 in tumor (in early or advanced stage or in both) were defined as tumor specific metabolism-immune gene pairs. For these pairs, the number of cancers which they existed in was counted to define the universality and specifity pairs. The pairs involved in only one cancer were defined as specifity. Otherwise, they were universality pairs. Diffrerentially expressed gene(DEG) analysis was performed using DEseq2^[35]^, raw counts of genes was used and pvalue was corrected by IHW method^[36]^. Genes with fold change greater than 1.5 and adjusted p value less than 0.05 was considered as DEGs.

### Functional analysis

An R package “clusterProfiler” was used to perform KEGG (Kyoto Encyclopedia of Genes and Genomes) and GO (Gene Ontology) analysis on the universality and specifity gens pairs^[37]^. The R package “heatmap” was used to draw the heatmap.

## Results

### Statistics of data

Totally, 5645 samples in 10 cancer groups were included in our research. Each group was divided into 7 groups: healthy control, tumor with clinical stage 1 to stage 4, early stage tumor (tumor clinical stage 1 and stage 2), advanced stage tumor (tumor clinical stage 3 and stage 4). For each group, correlation coefficient values were calculated between immune and metabolism related genes. Co-expressed pairs were further filtered into 3 groups: 0.7(|*r*| > 0.7), 0.8 (|*r*| > 0.8) and 0.9 (|*r*| > 0.9). The number of pairs in these groups were counted and presented in Figure 1, Figure 2 and Supplement Table 3. The number of co-expressed metabolism-immune pairs in HC is several to hundreds of times as many as that in tumors, except THCA. In THCA case, HC and tumor had the same order of magnitude of pairs number in |*r*| > 0.8 and |*r*| > 0.7 group, the number that in some tumor stages was even slightly more than that in HC. In addition, the co-expressed gene pairs show differences among cancers. In KIRC, KIRP and STAD, the co-expressed gene pair numbers was dozens of times greater than other cancers in quantity. Besides, the particular gene pairs were great different between HC and tumors.

**Figure 1.**
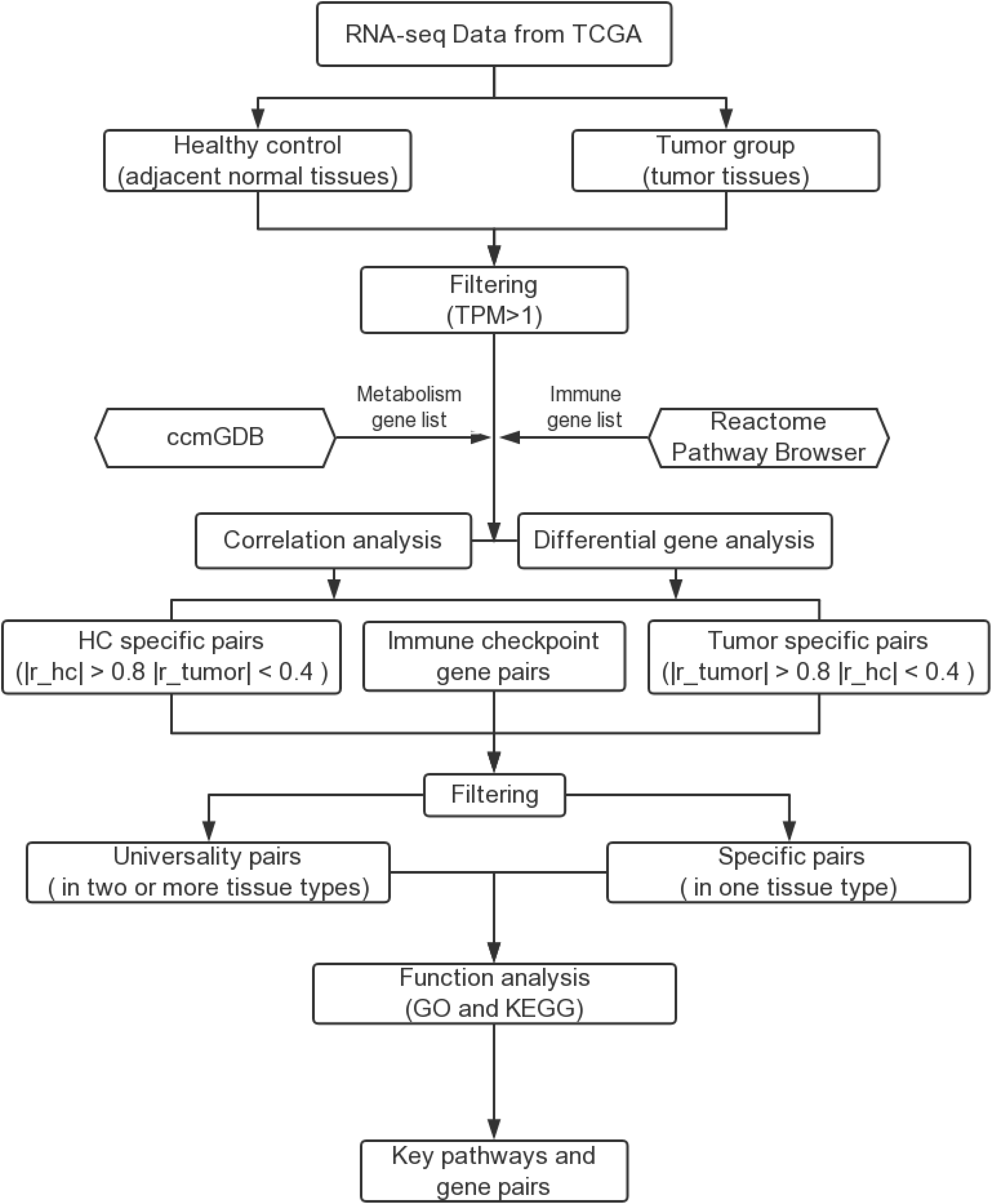
Flow chart of the integrated analysis of co-expressed pairs. The mRNA expression data were obtained from The Cancer Genome Atlas (TCGA). 5645 samples across 11 cancers were included in this research. According to the clinical information, each cancer was classified into 5 groups: healthy control (HC) and tumor samples from stage 1 to stage 4. For each gene with mean TPM less than 1 was filtered out. The list of human cellular metabolism genes was downloaded from ccmGDB. The immune-related gene list was obtained from the Reactome database. The spearman’s rank correlation was calculated between immune and metabolism genes and the differential gene analysis was analysed between HC and tumor samples. The correlation pairs with |r_hc| > 0.8 in HC and |r_tumor| < 0.4 were defined as HC specific metabolism-immune gene pairs. Pairs with |r_hc| < 0.4 in HC and |r_tumor| > 0.8 in tumor were defined as tumor specific pairs. The pairs including immune checkpoint were the immune checkpoint pairs. Besides, the pairs involved in only one cancer were defined as specific, otherwise, they were universal pairs. Then KEGG and GO enrichment analysis was performed between these pairs and the key pairs were screened out.

**Figure 2.**
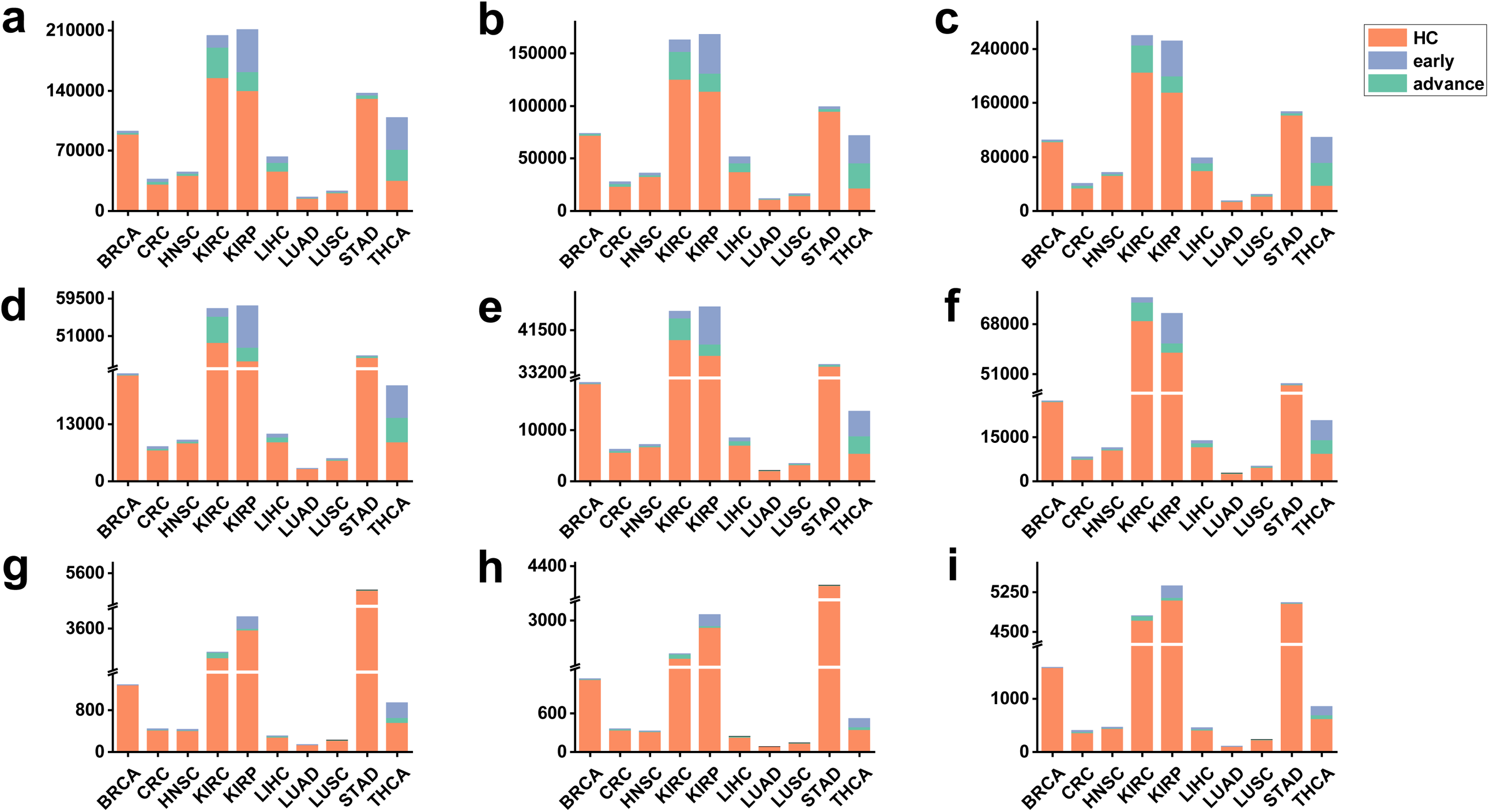
The quantitative comparison of co-expressed gene pairs between HC and tumor early/advanced stage tissues. Fig.abc, Fig.def and Fig.ghi represent comparison among HC, early and advanced stage in result of level0.7, level0.8 and level0.9 respectively. Fig.adg, Fig.beh and Fig.cfi represent comparison among HC, early and advanced stage in result of adaptive immunity, cytokine signaling pathway and innate immunity groups, respectively. The data of three levels all reflect the same rule that co-expressed gene pairs in HC are much more than those in early and advanced stages among all types of cancer, the proportions of HC, early and advanced stage are different among cancers.

### Universality and specifity of cancer types

Further filter was conducted to find out universal and specific metabolism-immune gene pairs across various types of cancer. The number of HC specific pairs was also much more than tumor specific pairs. The gene pairs were then combined with the number of their occurrences in cancer types. The pairs involved in only one cancer were defined as specific, otherwise, they were universal pairs. The full result is presented in Supplement Table 4-7. The top universal metabolism-immune gene pairs in HC and tumor were presented in Table 1 and Table 2, respectively. In HC group, several universal co-expressed gene pairs existed in more than 4 tissues, for example: *MED4-ACTR10, LSM5-ACTR10* and *NUPL2-UBA5*, etc. In the mean time, top universal co-expressed genes existed in more than 3 tumor tissues, these pairs are: *ADH1B-C7, GNG2-PTPRC, GNGT2-LAIR1, GNGT2-IL16, INHBA-COL3A1* and *CD38-PTPN7*.

### Functional analysis

In order to find out biological meanings of universal and tissue-specific metabolism-immune gene pairs, KEGG and GO enrichment analysis was performed in two system genes, respectively. Specific pairs of each cancer and universal pairs were separated to perform analysis, the grouping was similar with before: gene pairs were grouped according to 3 groups of immune genes, for each group, metabolism genes and immune genes were separated to perform KEGG and GO analysis. KEGG and GO terms derived from specifc pairs were presented in Suppliment Table 8-11. The function annotation of universal pairs were presented in Suppliment Table 12-15.

In KEGG and GO analysis, there were pathways which had a higher frequency of occurrence in cancer types, which means although different cancer types had differently co-expressed metabolism-immune gene pairs, these genes tended to show up in several same pathways. The pathways enriched to the most cancer types were screened out as the top KEGG and GO terms and shown in Supplement Table 16. There were also existed specific pathways indicating the pairs enriched in different pathways depending on the tumor types. Several metabolism pathways only showed in STAD and LUSC, respectively.

In the specific gene pairs group, the top KEGG pathways were phosphatidylinositol signaling system and inositol phosphate metabolism which appeared in 6 tumor types and in 10 normal tissues. Notably, the pathway including DGKs and PIP4Ks genes was only showed in normal types. In the universality gene pairs group, most genes strongly correlated with immune related genes were enriched in purine metabolism in normal tissues. Correspondingly, apelin signaling pathway including most metabolism genes was showed in tumor samples. The immune related genes were also analyzed by the functional enrichment and showed in Suppliment Table 12-15.

### The correlation between immune checkpoint and metabolism genes

To better understand the potential role of dysregulated gene pairs in immunotherapy, we choose the targetable immune checkpoint genes *TNFRSF4, CTLA4, PDCD1, CD274* and analysed the correlation between them and metabolism genes. We observed the tissue specificity and a consistent change that the correlation coefficient was much higher in normal than in tumor. Strong correlation between metabolism genes and immune checkpoint genes was showed in the one normal tissue or some HC of STAD, KIRP, KIRC and BRCA. Except for these four tissues, the correlation is not strong across other normal types(r > 0.8 as strong correlation). Looking at individual immune checkpoint genes, the most metabolism genes were correlated with *CD274* in the normal tissues of LIHC, KIRP, KIRC and BRCA. Only three genes were strongly correlated with *CTLA4* in HC of THCA and STAD. The genes correlated with *TNFRSF4* mainly enriched in normal tissues of STAD, the individuals were in HC of THCA(Figure 3A).

**Figure 3.**
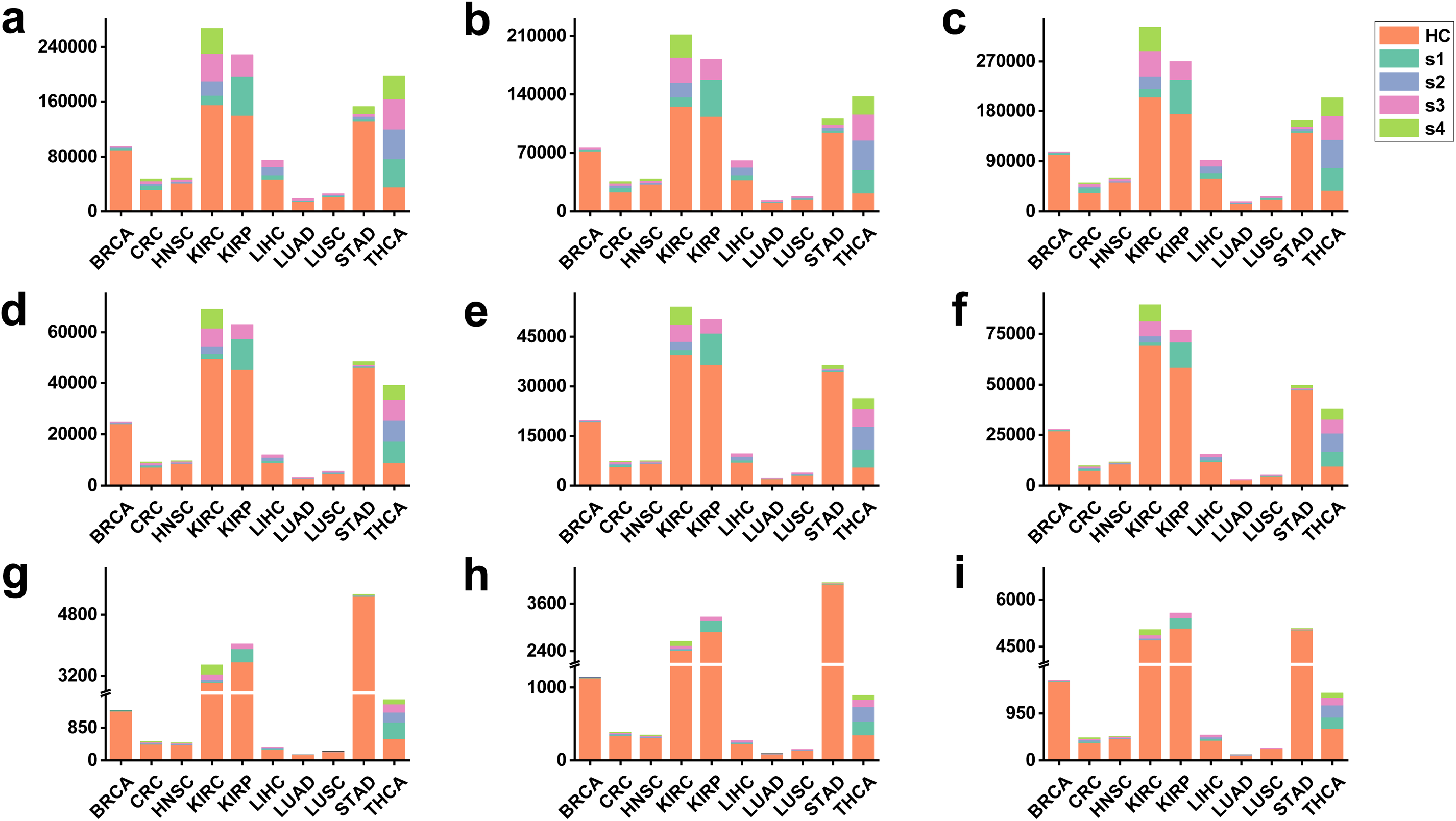
The quantitative comparison of co-expressed gene pairs between HC and tumor s1 to s4 stage tissues. The literal explanations also see in Figure 2.

**Figure 4.**
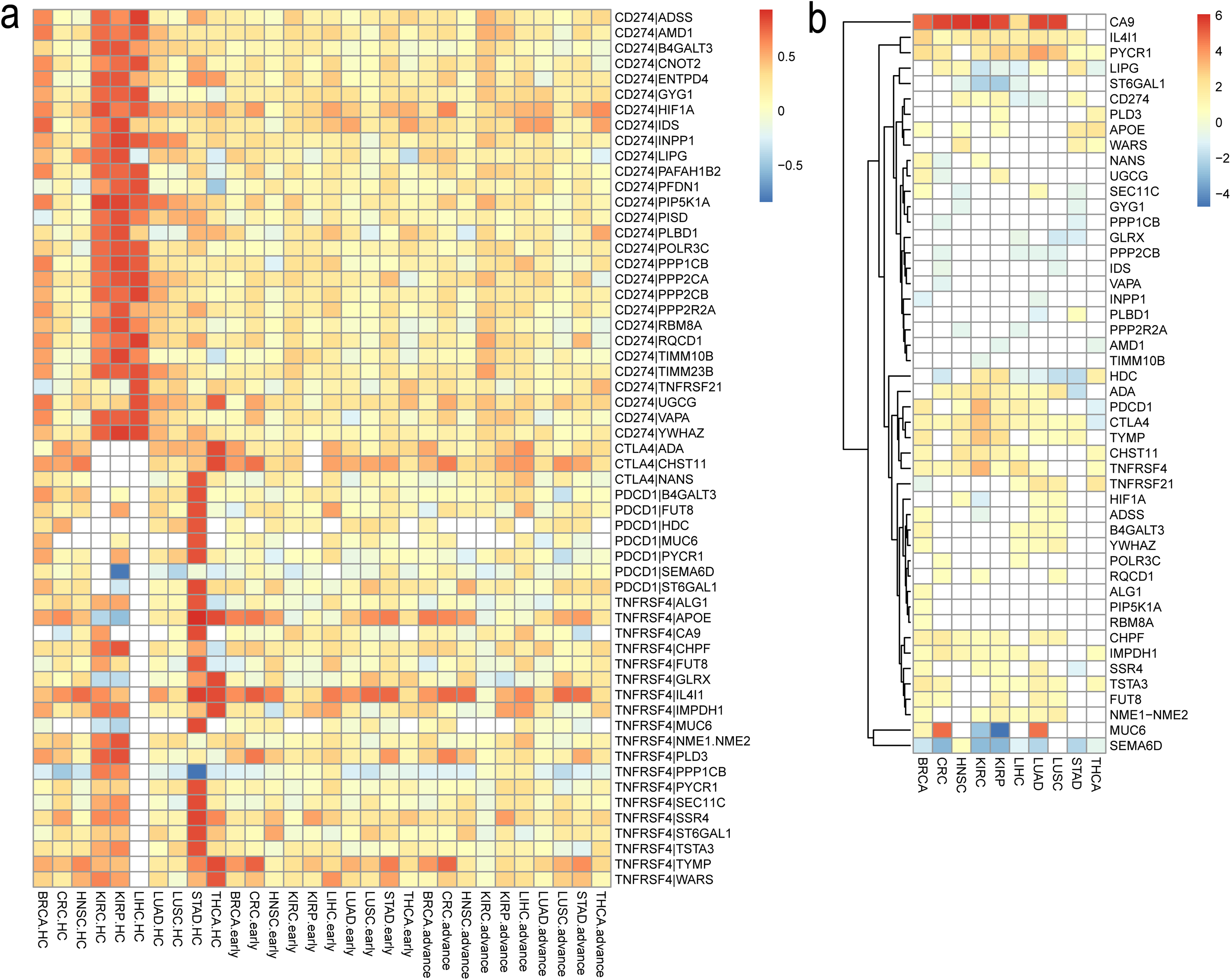
Hierarchical clustering of immune checkpoint gene pairs. Fig.a is the spearman correlation heatmap of metabolic genes with immune checkpoint genes. Red color illustrates a very strong positive correlation (r > 0.8), white represents missing value, and blue represents a negative correlation (r < −0.8). Fig.b represents the clustering of genes in the immune checkpoint gene pairs. Red color illustrates the gene was up-regulated in tumor compared with normal tissues(fold change > 1.5), white represents non-significant difference, blue represents gene was down-regulated (fold change < −1.5).

In order to rationalize the importance of above immune checkpoint related-genes in tumor, the differential expression levels were analysed. *CD274* was up-expressed in STAD, HNSC, KIRP and KIRC. *CTLA4* was up-regulated across cancer types expect THCA. *PDCD1* was up-regulated in 5 cancer types. In the metabolism genes, *ADA, CA9, CHPF, IL4I1* and *PYCR1* were all up-regulated at least 7 cancer types. The changed correlation may not mainly due to altered gene expression between normal and tumor, such as, *TNFRSF4* and *MUC6* was strongly correlated in normal tissues of STAD, differential expression of *TNFRSF4* or *MUC6* was not observed in STAD(Figure 3B).

## Discussion

Cancers are not just masses of malignant cells but including extracellular components and stromal cells, the malignant and these non-malignant cells and their cell-to-cell signalling creat the TME. However, the crosstalks between them is complicated and affected by a lot of factors. Our study was focused to elucidate the intertwined links between cell metabolism and tumor immunity through related-genes’ correlation analysis in human cancer. The results indicated that the disordered interaction occurred between metabolism and immunity in the TME at the overall level. Our study also revealed most relevant pairs and most significant signal pathway turbulences at the gene level in normal and cancer types. Besides, the metabolism genes strongly correlated with immune checkpoint genes were enumerated which may provide the potential therapeutic targets.

Many important stroma cells interact each other dynamically, with expressing metabolic and some immune genes, which have been reported to have critical roles in the immune response in cancer tissues^[38]^. The correlation among genes in tissue might reflect important biological relationship and variation in TME. The number of strongly correlated metabolism-immune gene pairs in normal tissues is 2-200 times than in tumor and even in tumor stage 1, revealing the disordered relation occurred in early stages of tumor and during the development of cancer. The TME turbulence may be the main reason leading the correlation changes of gene pairs, such as, cancer cells are frequently surrounded by hypoxic and acid microenvironment, to survive, they have evolved multiple adaptations and summoned other cells to help them^[39, 40]^. Besides much alteration has emerged in immune cells and immune responses as well^[41]^. Thus, it’s easy to understand the changes of their correlation between tumor and normal samples. The signatures of immune and metabolism vary depending of different cancer and tissues^[42]^, including macrophages quantity, overall lymphocyte infiltration, glycolysis and other biological characteristics. In particular types, we found that the number of gene pairs was varied across normal and tumor tissues. This may be a consequence of different environments in the different tissues.

The specific metabolic genes which strongly correlated with immune genes are enriched in some same pathways across both in normal and cancer tissues. The phosphatidylinositol signaling system and inositol phosphate metabolism have been identified in all normal tissues and 6 cancer types. The alteration of above pathways including gene expression, mutation, and posttranslational modification on key phospholipases and kinases are not only observed in tumor cells^[43, 44]^ but also in immune cells^[45, 46]^ and stroma cells^[47, 48]^, control the switch between immune stimulation and suppression in TME, which indicates phosphatidylinositol signaling system and inositol phosphate metabolism are closely relevant with immunoregulation in cancer. The metabolic genes in the two pathways which strongly correlated with immune genes in normal but missing in tumor may be the key gene connecting to immune disorder and tumor escape in tumor, such as, DGKs and PI5P4Ks family. DGKs family are considered as physiologic regulators of T- and NK-cell development and function through regulating TCR-signaling.

CD8-T cells and NK-cell activity can be enhanced and immune escape can be prevented through inhibition of DGKA in CRC^[49]^. *DGKA* was up-regulated in KIRP, KIRC and LUAD, in our research, *DGKA* may represent an key immunomodulator influencing the activity of immunity of immune cells in these TME. mTORC1 signaling influences the immune system by regulating diverse immune cell types, The enzyme encoded by *PIP4K2C* gene is a substrate of mTORC1^[50]^ and negatively regulates mTORC1, which indicate a close relationship between *PIP4K2C* and immune cells. Pip4k2c knockdown mice displayed increased immune infiltrates in various tissues, including liver, intestine, kidney, and lungs, these infiltrating cells are mostly T cells and B cells^[51]^. *PIP4K2C* was up-regulated in BRCA, which suggest *PIP4K2C* may influence the immune infiltrates in BRCA. Thus, the correlation analysis between metabolic genes and immune genes could identify some key metabolic genes which have impact on immune systems, however, how the other key metabolic genes (Supplement Table 17) influence the immune systems, we need further studies.

We observed the correlation was much higher in normal than in tumor between metabolism genes and immune checkpoint genes. The strongly related pairs were mainly showed in the HC of STAD, KIRP, KIRC and BRCA, and the specific correlation varied across normal tissues. These results suggest that correlation between immune checkpoints and metabolism genes is tissue specific and have disordered in tumors. We find a positive correlation between *HIF1A* and *PDL1* expression across cancers but not as strong as in normal tissues. *HIF1A* can regulate PDL1 expression at both the mRNA and protein level in myeloid-derived suppressor cells and also confirmed in melanoma and breast carcinoma bearing mice^[52]^. *HIF1A* was up-regulated in LUAD and LUSC from our data suggesting it may participate in *PDL1* regulation in these two cancers. Study shows that *FUT8* is the a post-translational regulator of *PDCD1* expression and blocking *FUT8* can enhance T cell activation under antigen presentation^[53]^. *FUT8* had a close relationship with *PDCD1* in HC of STAD while the correlation was decreased in tumors and it had a higher expression in LUAD, LUSC, CRC and BRCA, which indicates *FUT8* may influence immunity of T cell by affecting *PDCD1* expression in these tumor types. IL4I1 is the most described member of a family of immunosuppressive enzymes produced by antigen presenting cells which may act on T cells by direct inhibition of effector cell proliferation^[54]^. In this study, *IL4I1* is correlated with *TNFRSF4* in most cancer and normal types and up-regulated in all cancer types except STAD. Highly expressed *IL4I1* may play an important role in immune escape by affecting immune cell through *TNFRSF4*. Future research is needed to focus on the verification of these gene pairs associated with immune checkpoints.

## Conclusion

The disturbance of correlation between metabolism and immune related genes is revealed in our study. Phosphatidylinositol signaling system and inositol phosphate metabolism are strongly correlated with immune system both in normal and cancer tissues. The metabolic genes in the two pathways which strongly correlated with immune genes in normal but missing in tumor, such as DGKs and PI5P4Ks family, may be the key gene connecting to immune disorder and tumor escape in tumor. Particularly, the metabolic genes, such as, *HIF1A* and *FUT8* correlated with immune checkpoint are screened out which may be potential immunomodulator influencing therapeutic effect. The results including strongly correlated pairs and their pathways may open new avenues for exploration of the mechanisms underlying metabolism reprogramming and immune escape in TME.

## List of abbreviations

HNSC: Head and Neck Squamous Cell Carcinoma
KIRC: Kidney Renal Clear Cell Carcinoma
KIRP: Kidney Renal Papillary Cell Carcinoma
LIHC: Liver Hepatocellular Carcinoma
LUAD: Lung Adenocarcinoma
LUSC: Lung Squamous Cell Carcinoma
STAD: Stomach Adenocarcinoma
THCA: Thyroid Carcinoma.
BRCA: Breast Invasive Carcinoma
CRC: Colorectal Carcinoma
HC: Healthy Control
TCGA: The Cancer Genome Atlas
TME: Tumor Microenvironment
DGKs: Diacylglycerol Kinases
PIP4ks: Phosphatidylinositol-5-Phosphate 4-Kinases
FUT8: Fucosyltransferase 8
TNFRSF4: TNF Receptor Superfamily Member 4
CTLA4: Cytotoxic T-Lymphocyte Associated Protein 4
PDCD1: Programmed Cell Death 1
CD274: Programmed Cell Death Ligand 1
DGKA: Diacylglycerol Kinase Alpha
PIP4K2C: Phosphatidylinositol-5-Phosphate 4-Kinase Type 2 Gamma
HIF1A: Hypoxia Inducible Factor 1 Subunit Alpha
IL4I1: Interleukin 4 Induced 1

## Declarations

### Ethics approval and consent to participate

No ethical approval was required for this study.

### Consent for publication

Not applicable.

## Availability of data and material

Data of all considered in TCGA are publicly available at The Genomic Data Commons Data Portal (https://portal.gdc.cancer.gov/).

The datasets supporting the conclusions of this article are included within the article (and its additional files).

## Authors’ contributions

Hongli Du designed and conceived study. Zixi Chen conducted analyses. Jinfen Wei and Zixi Chen wrote the manuscript. Yuchen Yuan and Ying Cui assisted with conducting analyses the data. Yanyu Zhang and Yumin Hu offered suggestions for the analysis and manuscript writing. Hongli Du were involved in revising the manuscript. All authors read and approved the final manuscript.

## Acknowledgements

This study would have been impossible without the comprehensive data sets made publicly available by the TCGA Research Network.

## Additional files

Additional file 1 - Sample size of each group.xlsx

Additional file 2 - Gene list of four groups(metabolism, innate immunity, adaptive immunity, cytokine signaling pathways).xlsx

Additional file 3 - Full table of statistics(The number of pairs with |r| > 0.9,0.8,0.7 in each group across cancer types).xlsx

Additional file 4 - Universalilty pairs in HC group(the pairs in at least 2 normal tissues).xlsx

Additional file 5 - Specifity pairs in HC group(the pairs in only 1 normal tissue).xlsx

Additional file 6 - Universalilty pairs in tumor group(the pairs in at least 2 cancer types).xlsx

Additional file 7 - Specifity pairs in tumor group(the pairs in only 1 cancer type).xlsx

Additional file 8 - GO terms derived from specifity pairs in HC.xlsx

Additional file 9 - KEGG terms derived from specifity pairs in HC.xlsx

Additional file 10 - GO terms derived from specifity pairs in tumor.xlsx

Additional file 11 - KEGG terms derived from specifity pairs in tumor.xlsx

Additional file 12 - GO terms derived from universality pairs in HC.xlsx

Additional file 13 - KEGG terms derived from universality pairs in HC.xlsx

Additional file 14 - GO terms derived from universality pairs in tumor.xlsx

Additional file 15 - KEGG terms derived from universality pairs in tumor.xlsx

Additional file 16 - Top pathways counts(the pathways in most tissues and cancer types).xlsx

Additional file 17 - The genes involved in phosphatidylinositol signaling system and inositol phosphate metabolism across normal and cancer types.xlsx

## References

[1] Martinez-Bosch N, Vinaixa J, Navarro P. Immune Evasion in Pancreatic Cancer: From Mechanisms to Therapy. Cancers (Basel). 2018. 10(1).

[2] Muenst S, Läubli H, Soysal SD, Zippelius A, Tzankov A, Hoeller S. The immune system and cancer evasion strategies: therapeutic concepts. J Intern Med. 2016. 279(6): 541–62.

[3] Panda AK, Bose S, Sarkar T, et al. Cancer-immune therapy: restoration of immune response in cancer by immune cell modulation. Nucleus. 2017. 60(2): 1–17.

[4] Boon T. Teaching the immune system to fight cancer. Sci Am. 1993. 268(3): 82.

[5] Gubin MM, Zhang X, Schuster H, et al. Checkpoint blockade cancer immunotherapy targets tumour-specific mutant antigens. Nature. 2014. 515(7528): 577–581.

[6] Burugu S, Dancsok AR, Nielsen TO. Emerging targets in cancer immunotherapy. Semin Cancer Biol. 2017. 52(Pt 2): S1044579X17301827.

[7] Mayes PA, Hance KW, Hoos A. The promise and challenges of immune agonist antibody development in cancer. Nat Rev Drug Discov. 2018. 17(7).

[8] Gang C, Huang AC, Wei Z, et al. Exosomal PD-L1 contributes to immunosuppression and is associated with anti-PD-1 response. Nature. 2018.

[9] Yoshida GJ. Metabolic reprogramming: the emerging concept and associated therapeutic strategies. Journal of Experimental & Clinical Cancer Research Cr. 2015. 34(1): 1–10.

[10] Vander Heiden MG, Cantley LC, Thompson CB. Understanding the Warburg Effect: The Metabolic Requirements of Cell Proliferation. Science. 2009.

[11] Liberti MV, Locasale JW. The Warburg Effect: How Does it Benefit Cancer Cells. Trends Biochem Sci. 2016. 41(3): 211–218.

[12] Mills EL, Kelly B, O&Aposneill LAJ. Mitochondria are the powerhouses of immunity. Nat Immunol. 2017. 18(5): 488.

[13] Renner K, Singer K, Koehl GE, et al. Metabolic Hallmarks of Tumor and Immune Cells in the Tumor Microenvironment. Front Immunol. 2017. 8(6): 248.

[14] Ma C, Han M, Heinrich B, et al. Gut microbiome-mediated bile acid metabolism regulates liver cancer via NKT cells. Science. 2018. 360(6391): eaan5931.

[15] Subhra&nbsp, Biswas K. Metabolic Reprogramming of Immune Cells in Cancer Progression. Immunity. 2015. 43(3): 435–449.

[16] Porta C, Sica A, Riboldi E. Tumor-associated myeloid cells: new understandings on their metabolic regulation and their influence in cancer immunotherapy. FEBS J. 2017. 285(4): 717–733.

[17] Lakins MA, Ghorani E, Munir H, Martins CP, Shields JD. Cancer-associated fibroblasts induce antigen-specific deletion of CD8 + T Cells to protect tumour cells. Nat Commun. 2018. 9(1).

[18] Bieniasz M, Oszajca K, Eusebio M, Kordiak J, Bartkowiak J, Szemraj J. The positive correlation between gene expression of the two angiogenic factors: VEGF and BMP-2 in lung cancer patients. Lung Cancer. 2009. 66(3): 319–326.

[19] Punt S, Houwing-Duistermaat JJ, Schulkens IA, et al. Correlations between immune response and vascularization qRT-PCR gene expression clusters in squamous cervical cancer. Mol Cancer. 2015. 14(1): 71.

[20] Bindea G, Mlecnik B, Tosolini M, et al. Spatiotemporal dynamics of intratumoral immune cells reveal the immune landscape in human cancer. Immunity. 2013. 39(4): 782–795.

[21] Hu J, Locasale JW, Bielas JH, et al. Heterogeneity of tumor-induced gene expression changes in the human metabolicnetwork. Nat Biotechnol. 2013. 31(6): 522–529.

[22] Gaude E, Frezza C. Tissue-specific and convergent metabolic transformation of cancer correlates with metastatic potential and patient survival. Nat Commun. 2016. 7: 13041.

[23] Li B, Severson E, Pignon JC, et al. Comprehensive analyses of tumor immunity: implications for cancer immunotherapy. Genome Biol. 2016. 17(1): 174.

[24] Thorsson V, Gibbs DL, Brown SD, et al. The Immune Landscape of Cancer. Immunity. 2018. 81(1): 105.

[25] Grossman RL, Heath AP, Ferretti V, et al. Toward a Shared Vision for Cancer Genomic Data. N Engl J Med. 2016. 375(12): 1109.

[26] Kim P, Cheng F, Zhao J, Zhao Z. ccmGDB: a database for cancer cell metabolism genes. Nucleic Acids Res. 2016. 44(Database issue): D959–D968.

[27] Ikezu T. Innate Immunity Signaling. 2017.

[28] Fabregat A, Sidiropoulos K, Garapati P, et al. The Reactome pathway Knowledgebase. Nucleic Acids Res. 2014. 42(Database issue): 472–7.

[29] Dillies MA, Rau A, Aubert J, et al. A comprehensive evaluation of normalization methods for Illumina high-throughput RNA sequencing data analysis. Brief Bioinform. 2013. 14(6): 671–683.

[30] Wagner GP, Kin K, Lynch VJ. Measurement of mRNA abundance using RNA-seq data: RPKM measure is inconsistent among samples. Theory Biosci. 2012. 131(4): 281–285.

[31] Mandric I, Temate-Tiagueu Y, Shcheglova T, Al SS, Zelikovsky A, Mandoiu II. Fast Bootstrapping-Based Estimation of Confidence Intervals of Expression Levels and Differential Expression from RNA-Seq Data. Bioinformatics. 2017. 33(20).

[32] Harrow J, Denoeud F, Frankish A, et al. GENCODE: producing a reference annotation for ENCODE. Genome Biol. 2006. 7(Suppl 1): 1–9.

[33] Spearman C. The proof and measurement of association between two things. By C. Spearman, 1904. American Journal of Psychology. 1987. 100(3/4): 441–471.

[34] Sakurai T, Kondoh N, Arai M, et al. Functional roles of Fli-1, a member of the Ets family of transcription factors, in human breast malignancy. Cancer Sci. 2010. 98(11): 1775–1784.

[35] Love MI, Huber W, Anders S. Moderated estimation of fold change and dispersion for RNA-seq data with DESeq2. Genome Biol. 2014. 15(12): 550.

[36] Ignatiadis N, Klaus B, Zaugg JB, Huber W. Data-driven hypothesis weighting increases detection power in genome-scale multiple testing. Nat Methods. 2016. 13(7): 577–80.

[37] Yu G, Wang LG, Han Y, He QY. clusterProfiler: an R package for comparing biological themes among gene clusters. OMICS. 2012. 16(5): 284–287.

[38] Lim B, Woodward WA, Wang X, Reuben JM, Ueno NT. Inflammatory breast cancer biology: the tumour microenvironment is key. Nat Rev Cancer. 2018. 18(8): 485–499.

[39] Huber V, Camisaschi C, Berzi A, et al. Cancer acidity: An ultimate frontier of tumor immune escape and a novel target of immunomodulation. Semin Cancer Biol. 2017. 43: 74–89.

[40] Terry S, Buart S, Chouaib S. Hypoxic Stress-Induced Tumor and Immune Plasticity, Suppression, and Impact on Tumor Heterogeneity. Front Immunol. 2017. 8: 1625.

[41] Correction: Bruton Tyrosine Kinase-Dependent Immune Cell Cross-talk Drives Pancreas Cancer. Cancer Discov. 2016. 6(7): 802.

[42] Gaude E, Frezza C. Tissue-specific and convergent metabolic transformation of cancer correlates with metastatic potential and patient survival. Nat Commun. 2016. 7: 13041.

[43] Yousef AI, El-Masry OS, Abdel Mohsen MA. Impact of Cellular Genetic Make-up on Colorectal Cancer Cell Lines Response to Ellagic Acid: Implications of Small Interfering RNA. Asian Pac J Cancer Prev. 2016. 17(2): 743–8.

[44] Chen H, Gao J, Du Z, Zhang X, Yang F, Gao W. Expression of factors and key components associated with the PI3K signaling pathway in colon cancer. Oncol Lett. 2018. 15(4): 5465–5472.

[45] Best SA, De Souza DP, Kersbergen A, et al. Synergy between the KEAP1/NRF2 and PI3K Pathways Drives Non-Small-Cell Lung Cancer with an Altered Immune Microenvironment. Cell Metab. 2018. 27(4): 935-943.e4.

[46] Elich M, Sauer K. Regulation of Hematopoietic Cell Development and Function Through Phosphoinositides. Front Immunol. 2018. 9: 931.

[47] Cho Y, Cho EJ, Lee JH, et al. Hypoxia Enhances Tumor-Stroma Crosstalk that Drives the Progression of Hepatocellular Carcinoma. Dig Dis Sci. 2016. 61(9): 2568–77.

[48] Hirsch E, Ciraolo E, Franco I, Ghigo A, Martini M. PI3K in cancer-stroma interactions: bad in seed and ugly in soil. Oncogene. 2014. 33(24): 3083–90.

[49] Noessner E. DGK-α: A Checkpoint in Cancer-Mediated Immuno-Inhibition and Target for Immunotherapy. Front Cell Dev Biol. 2017. 5: 16.

[50] Mackey AM, Sarkes DA, Bettencourt I, Asara JM, Rameh LE. PIP4kγ is a substrate for mTORC1 that maintains basal mTORC1 signaling during starvation. Sci Signal. 2014. 7(350): ra104.

[51] Shim H, Wu C, Ramsamooj S, et al. Deletion of the gene Pip4k2c, a novel phosphatidylinositol kinase, results in hyperactivation of the immune system. Proc Natl Acad Sci U S A. 2016. 113(27): 7596–601.

[52] Noman MZ, Desantis G, Janji B, et al. PD-L1 is a novel direct target of HIF-1α, and its blockade under hypoxia enhanced MDSC-mediated T cell activation. J Exp Med. 2014. 211(5): 781–90.

[53] Okada M, Chikuma S, Kondo T, et al. Blockage of Core Fucosylation Reduces Cell-Surface Expression of PD-1 and Promotes Anti-tumor Immune Responses of T Cells. Cell Rep. 2017. 20(5): 1017–1028.

[54] Cousin C, Aubatin A, Le Gouvello S, Apetoh L, Castellano F, Molinier-Frenkel V. The immunosuppressive enzyme IL4I1 promotes FoxP3(+) regulatory T lymphocyte differentiation. Eur J Immunol. 2015. 45(6): 1772–82.

